# COUP-TFII regulates early bipotential gonad signaling and commitment to ovarian progenitors

**DOI:** 10.1101/2023.08.09.552582

**Authors:** Lucas G. A. Ferreira, Marina M. L. Kizys, Gabriel A. C. Gama, Svenja Pachernegg, Gorjana Robevska, Andrew H. Sinclair, Katie L. Ayers, Magnus R. Dias da Silva

## Abstract

The absence of expression of the Y-chromosome linked testis-determining gene *SRY* in early supporting gonadal cells (ESGC) of bipotential gonads leads to ovarian development. However, genetic variants in *NR2F2*/COUP-TFII represent a novel cause of *SRY*-negative 46,XX testicular/ovotesticular differences of sex development (T/OT-DSD). Thus, we hypothesized that COUP-TFII is part of the ovarian developmental network. We examined *NR2F2*/COUP-TFII expression and the genetic network under its regulation in human gonadal cells by analyzing single cell RNA-sequencing datasets of fetal gonads, differentiating induced pluripotent stem cells into bipotential gonad-like cells *in vitro*, and generating a *NR2F2* knockout (KO) in the human granulosa-like cell line COV434. *NR2F2* expression is highly upregulated during the bipotential gonad development, being detected in ESGCs. We identified that *NR2F2* ablation in COV434 cells downregulated markers of ESGC and pre-granulosa cells, suggesting that COUP-TFII has a role in maintaining a multipotent state necessary for commitment to the ovarian development. We propose that impairment of COUP-TFII function may disrupt the transcriptional plasticity of ESGCs and instead drive them into commitment to the testicular pathway.

## Introduction

The bipotential gonad is an embryonic tissue harboring multipotent cells with the capacity to adopt either a testicular or an ovarian cell fate. From the end of the sixth week of human gestation, the early supporting gonadal cells (ESGC) expressing the Y-chromosome linked testis-determining gene *SRY* give rise to fetal Sertoli cells expressing a genetic network that promotes testis development and antagonizes ovary development. Conversely, the *SRY*-negative ESGCs develop as pre-granulosa cells by repressing testis determination (Reyes *et al*, 2023). However, the identification of individuals with an *SRY*-negative 46,XX karyotype and testicular tissue that produces androgens suggests either upregulation of other testis genes or down regulation of ovarian genes (Syryn *et al*, 2023). These rare conditions are classified as 46,XX testicular or ovotesticular differences of sex development (T/OT-DSD) depending on the identification of gonads that resemble testes or the presence of both testicular and ovarian tissue (Grinspon & Rey, 2019). Although many genetic factors have been found to be involved in testis development, far less is known about genes regulating the formation of an ovary (Eggers *et al*, 2014; Stévant *et al*, 2019).

The translocation of the *SRY* gene is a common etiology of 46,XX T/OT-DSD (Berglund *et al*, 2017); however, our group and others have found that loss-of-function genetic variants in *NR2F2*, encoding the transcription factor COUP-TFII, represent a novel cause of *SRY*-negative cases (Carvalheira *et al*, 2019; Bashamboo *et al*, 2018). We previously described an individual with *SRY*-negative 46,XX OT-DSD who was born with atypical male external genitalia, cardiac defects, and blepharophimosis-ptosis-epicanthus inversus syndrome (BPES) (Carvalheira *et al*, 2019). The molecular investigation identified a *de novo* heterozygous 3-Mb deletion at 15q26.2 encompassing the *NR2F2* gene. Additionally, heterozygous frameshift variants in the *NR2F2* gene were identified in children presenting with *SRY*-negative 46,XX T/OT-DSD, virilized genitalia, congenital heart disease, and BPES (Bashamboo *et al*, 2018; Lambert *et al*, 2021). The mechanisms responsible for testis development in individuals with *NR2F2* genetic variants are yet to be understood.

COUP-TFII participates in the regulation of organogenesis, neuronal development, angiogenesis, cardiovascular development, reproduction, and metabolic processes (Polvani *et al*, 2020). During heart and vascular development, COUP-TFII is a major regulator of epithelial to mesenchymal transition and has been implicated in the induction of vein and lymphatic vessel identity and the repression of artery-specific genes (Polvani *et al*, 2020). Several variants in the *NR2F2* gene have indeed been reported in association with cardiac defects in humans (Al Turki *et al*, 2014). In mice, *Nr2f2* expression is detected in the visceral mesoderm and developing heart from embryonic day (e) 8.5 (Pereira *et al*, 1999). Homozygous deletion of *Nr2f2* is lethal around e10 due to cardiac defects, and two-thirds of the heterozygous *Nr2f2* mice die before puberty (Lin *et al*, 2012), indicating that COUP-TFII function during cardiovascular development is vital.

COUP-TFII also plays a role in the development of gonadal steroidogenic cells in males and females from different species. Complete knockout of *Nr2f2* in male mice at e18.5 and at pre-pubertal stages causes the development of dysfunctional adult Leydig cells (Qin *et al*, 2008). COUP-TFII is a marker of stem cells that give rise to the adult Leydig cell population and regulates *Star, Insl3*, and *Amhr2* gene expression by directly binding to their respective promoter sequences (Chen *et al*, 2012; de Mattos *et al*, 2022). Thus, COUP-TFII participates in the commitment of the progenitor cells into fully functional steroidogenic adult Leydig cells. In female mice, *Nr2f2* heterozygous deletion impairs sex steroid hormone synthesis in the ovary and decidualization in the uterus (Takamoto *et al*, 2005; Petit *et al*, 2007). Additionally, COUP-TFII has been implicated in Wolffian duct degradation in females (Zhao *et al*, 2017). However, these studies reported that both ovary signaling and morphology in mice with *Nr2f2* haploinsufficiency are mostly unperturbed, suggesting that COUP-TFII function during gonadal development may differ between humans and mice. Differences in transcript variants expressed by the *NR2F2* gene may contribute to divergencies between species. While the murine *Nr2f2* gene is transcribed in two mRNA variants, the human *NR2F2* expresses four variants (v1-4) from independent transcription start sites selecting alternative exon 1. These human transcript variants are translated into three distinct protein isoforms (Polvani *et al*, 2020).

*NR2F2* gene expression has been used as a marker of gonadal interstitial/mesenchymal progenitor cells (Rastetter *et al*, 2014; Smith *et al*, 2014). However, *NR2F2* expression has been detected in humans and mice prior to supporting and interstitial cell lineage differentiation. Male and female early somatic cells (the progenitors of supporting and interstitial cells) express *NR2F2* from six weeks of human gestation and e10.5 in mice (Stévant *et al*, 2019; Guo *et al*, 2021). We aimed to examine the expression of *NR2F2* transcript variants during early gonadal development in humans and to assess the genetic networks regulated by COUP-TFII in a gonadal cell context. Our data associates *NR2F2*/COUP-TFII activity with the maintenance of the early bipotential gonad state and with the commitment of ESGCs into female gonadal progenitors. Together our data suggest that ovary development is an active process and that COUP-TFII plays a central role in this cellular decision.

## Results

### NR2F2 expression is detected in progenitor cells of the human bipotential gonad and shifts to a sex-specific pattern during the commitment of supporting cells

To study *NR2F2* expression during early gonadal development, we examined single-cell RNA sequencing (scRNA-seq) datasets of somatic cell lineages from male and female human gonads between 6- and 21-weeks gestation (Garcia-Alonso *et al*, 2022). As shown in Fig 1A, *NR2F2*-expressing cells were identified in clusters related to the first wave of supporting cells, i.e. early somatic cells (cluster 2a), bipotential early supporting gonad cells (ESGC, cluster 2b), the first wave of pre-granulosa cells (preGC-I, cluster 2c), and early supporting *PAX8*-expressing cells (cluster 3a). The early somatic cells give rise to the ESGCs, which are bipotential precursors that give rise to the sex-specific supporting cell lineages in the early gonad. The early *PAX8*-expressing cells are sexually undifferentiated cells at the gonadal– mesonephric interface. We also detected the expected *NR2F2* expression in interstitial/mesenchymal cells, with the highest expression levels in gonadal interstitial cells (cluster 4a). *GATA2*-expressing coelomic epithelial cells (cluster 1a), which are associated with the extragonadal mesonephros, were also enriched for *NR2F2* expression. Conversely, the *GATA4*-expressing coelomic epithelial cells (cluster 1b), which give rise to early somatic cells and bipotent ESGCs, and the fetal Sertoli cells (cluster 2d) demonstrated low levels of *NR2F2* expression (Fig 1A, Appendix Fig S1).

**Fig 1.**
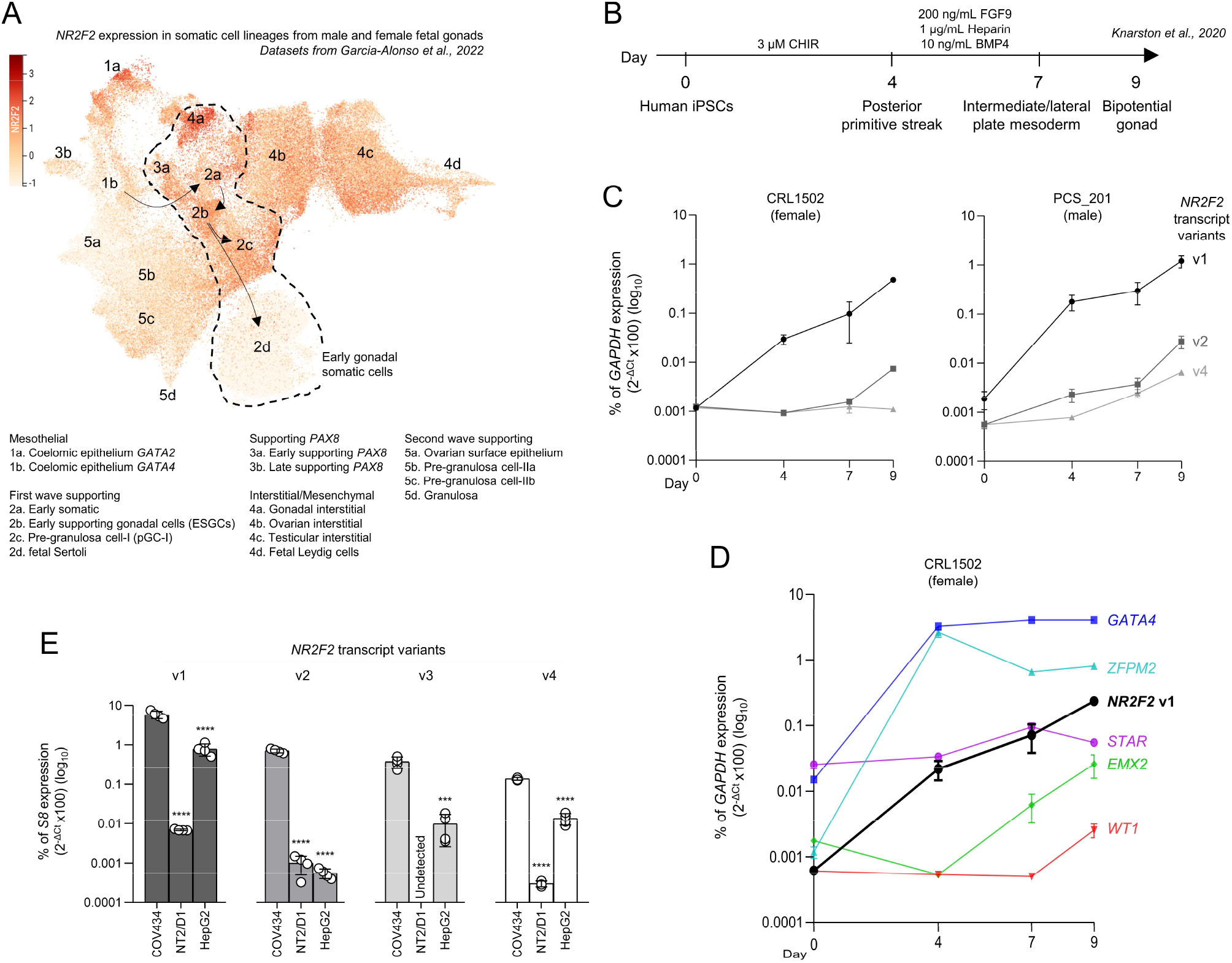
*NR2F2* expression during early human gonadal development and in COV434 and NT2/D1 cell lines. (**A**) The expression pattern of *NR2F2* projected on the UMAP plot showing cell lineages in the scRNA-seq datasets of male and female somatic cells obtained from human gonads (Garcia-Alonso *et al*, 2022). The color scale represents *NR2F2* gene expression levels. Arrows indicate the predicted developmental trajectory of cell lineages based on evidence from the literature. (**B**) The monolayer differentiation protocol for human embryonic bipotential gonad described by Knarston et al. (Knarston *et al*, 2020). (**C**,**D**) RT-qPCR data of gene expression of *NR2F2* transcript variants in CRL1502 (female) and PCS_201 (male) iPSCs (C), and of the bipotential gonad markers *GATA4, ZFPM2, EMX2*, and *WT1*, and the steroidogenic cell marker *STAR* (D) during the differentiation protocol. *NR2F2* v3 was not detected under these experimental conditions. Data were normalized as a percentage of *GAPDH* (reference gene) expression. Mean ± SEM (n=3). (**E**) RT-qPCR assay comparing the expression of *NR2F2* transcript variants (v1-4) in COV434, NT2/D1, and HepG2 cell lines. Data is represented as a percentage of *S8* (reference gene) expression. Mean ± SEM (n=4). One-Way ANOVA followed by Tukey test (v1, v2, and v4) and t-test (v3). Asterisks indicate statistical significance in relation to COV434 expression. ***p<0.001, ****p<0.0001.

Using the protocol previously established by Knarston et al. (Knarston *et al*, 2020) (Fig 1B), we examined the expression of *NR2F2* transcript variants during the differentiation of human female and male induced pluripotent stem cells (iPSC) through primitive streak, mesoderm, and bipotential gonad stages. The *NR2F2* transcript v1 was gradually upregulated during the differentiation of female and male iPSCs, presenting a nearly 1000-fold increase in bipotential gonad state at day 9 of culture when comparing to day 0 iPSCs. The *NR2F2* v2 and v4 were upregulated, but presented lower fold changes than v1, and the *NR2F2* v3 was not detected during the bipotential gonad differentiation in either cell line (Fig 1C). During this *in vitro* differentiation, *NR2F2* expression was upregulated along with bipotential gonad markers *GATA4, ZFPM2, EMX2*, and *WT1*, while the steroidogenic cell marker *STAR* was not upregulated, as shown for the female iPSCs (Fig 1D).

The analysis of scRNA-seq datasets of developing human gonads suggested that the differentiation of ESGCs into fetal Sertoli cells occurs in parallel with the loss of *NR2F2* expression, while preGC-I still expressed *NR2F2* (Fig 1A). In line with this observation, we found that all *NR2F2* transcript variants had higher expression levels in the granulosa-like cell line COV434 compared to the Sertoli-like cell line NT2/D1 and to male hepatocyte HepG2, an extragonadal cell line known to express *NR2F2* (Erdős & Bálint, 2019) (Fig 1E). We detected nearly 1000-fold higher expression of v1, v2, and v4 in COV434 cells than in NT2/D1 cells. The v3 was not detected in the NT2/D1 cell line. The v1, encoding the canonical protein isoform A, showed the highest abundance among the *NR2F2* transcript variants in all investigated cell lines (Fig 1E). This data indicates that COUP-TFII is an important transcription factor in the granulosa-like COV434 cell line.

### Generation and validation of a COV434 NR2F2-knockout cell line

To study the transcriptional networks regulated by COUP-TFII in a granulosa-like cell context, we generated COV434 cells with a mutation affecting all *NR2F2* transcript variants using CRISPR/Cas9. We used a guide RNA (gRNA) directed toward the 5’ region of exon 2, which is a shared exon among all the *NR2F2* transcripts (Appendix Fig S2, Fig 2A). After allele isolation and genotype screening by Sanger sequencing of *NR2F2* exon 2, we identified a potential wild-type (WT) COV434 clone and a clone with the homozygous mutation c.484delG (NM_021005) at the expected site. This mutation was predicted to generate a frameshift and a premature stop codon in the *NR2F2* gene (p.Gln163fs*4, NP_066285) (Fig 2B). Genotyping array confirmed that genome integrity was maintained (absence of aneuploidies or large copy number variations (>0.50 Mb)) in these COV434 clones (data not shown). Western blotting and RT-qPCR assays demonstrated the complete loss of COUP-TFII isoform A, possibly caused by nonsense-mediated mRNA decay, and the decrease in expression of the four *NR2F2* transcripts in the COV434 clone harboring the mutation c.484delG when compared to the WT clone and the non-transfected (NT) COV434 cells (Fig 2C,D). As functional validation, we examined the expression of known COUP-TFII target genes in endothelial cells using qPCR. *HEY2* and *E2F1*, but not *NOTCH1*, were downregulated in the *NR2F2*-KO *versus* the WT COV434 cell clone (Fig 2E). We therefore concluded that we had created a homozygous *NR2F2*-KO COV434 cell line, and moved on to further investigate the role of *NR2F2* using this cell line.

**Fig 2.**
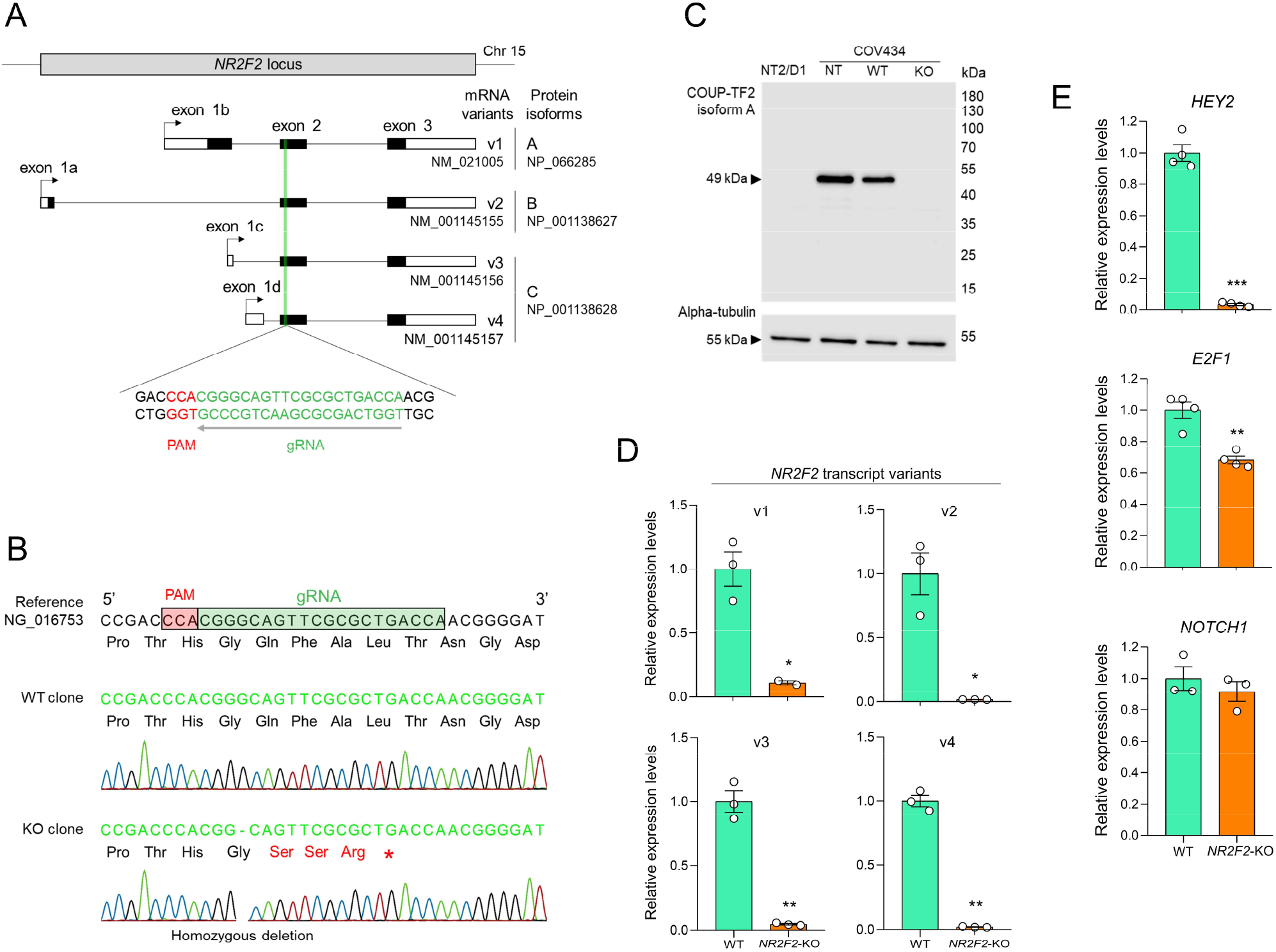
Generation and validation of a COV434 cell line carrying a knockout of the *NR2F2* gene. (**A**) Schematic representation of the human *NR2F2* locus on chromosome 15. Four mRNA variants generated by alternative transcription start sites, which encode three protein isoforms, are depicted. Filled boxes indicate coding sequences (CDS), empty boxes indicate untranslated regions and lines represent introns. Arrows indicate transcriptional start sites. The position of the guide RNA (gRNA) targeting exon 2 is indicated. PAM, protospacer adjacent motif. (**B**) Genotyping of representative isolated alleles from two selected single-cell clones by Sanger sequencing revealed potential wild-type (WT) and *NR2F2*-knockout (KO) clones. The KO clone presented the homozygous mutation c.484delG (NM_021005), p.Gln163fs*4 (NP_066285). (**C**) Western blotting assays using total protein extracts from NT2/D1, COV434 non-transfected (NT), WT, and *NR2F2*-KO cell clones. The anti-COUP-TFII used recognizes the human isoform A, expressed by the *NR2F2* variant 1. Alpha-tubulin was used as endogenous control. The molecular weight (kDa) is indicated. (**D, E**) RT-qPCR assay comparing relative mRNA expression of the four *NR2F2* transcript variants (D) and known COUP-TFII regulated genes (E) between WT and *NR2F2*-KO COV434 cell clones. *S8* was used as a reference gene. Values are mean ± SEM (n=3-4). Student’s t-test with Welch’s correction, *p<0.05, **p<0.01, ***p<0.001.

COV434 cells grow as diffuse sheets forming occasional follicle-like structures containing eosinophilic luminal material (van den Berg-Bakker *et al*, 1993). No difference in growth pattern or cytomorphology was observed between the WT and the *NR2F2*-KO COV434 cells when examined by H&E and staining of cytoskeletal proteins (Fig 3A,B).

**Fig 3.**
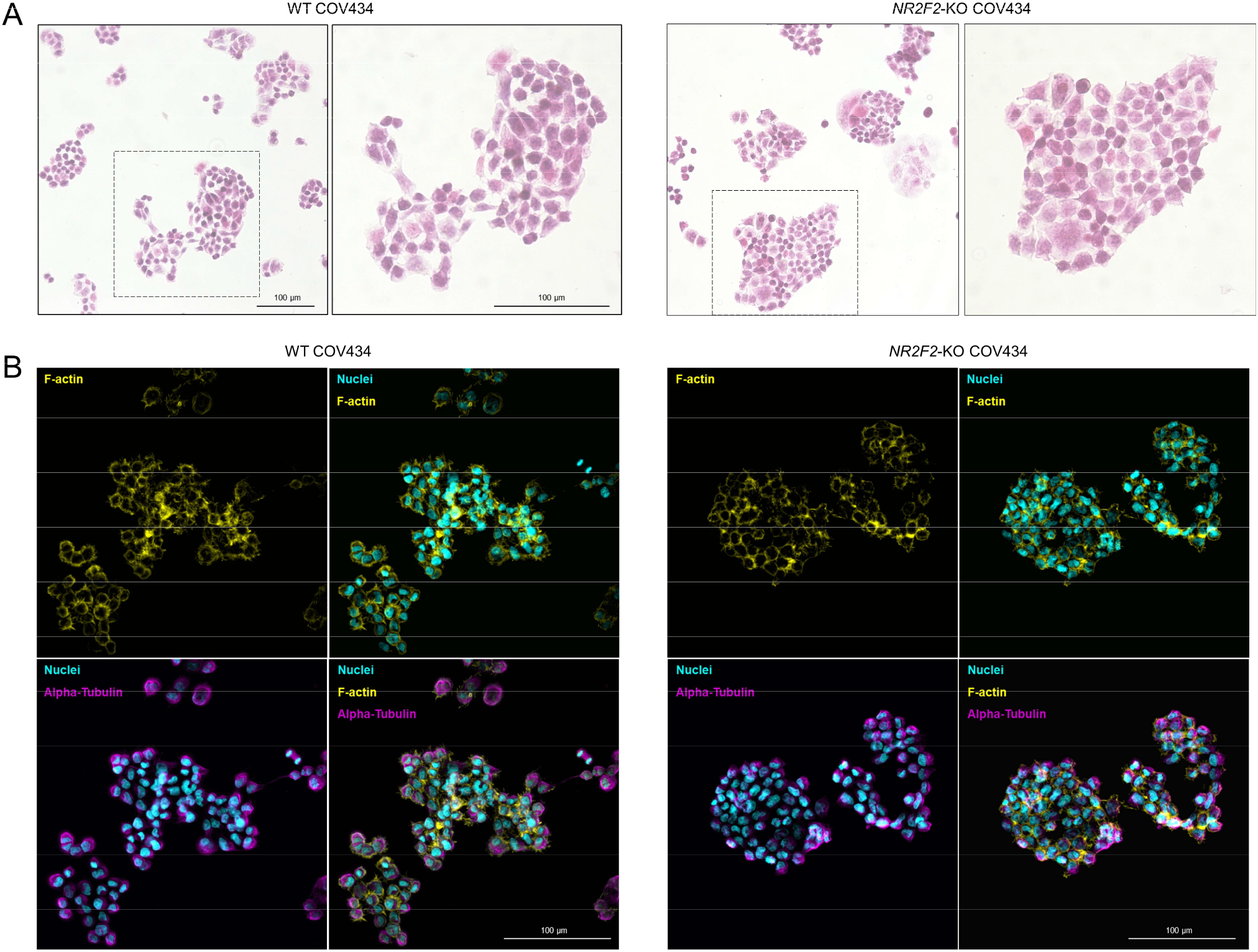
Cytomorphology examination of WT and the *NR2F2*-KO COV434 cells. (**A**) Light microscopy images of WT and *NR2F2*-KO COV434 cell clones stained with H&E. Magnifications of the areas outlined by the dashed boxes are shown on the right panels. (**B**) Fluorescence images of cells stained with phalloidin (yellow), DAPI (cyan), and Anti-Alpha-Tubulin (magenta). Merged images are shown.

### COUP-TFII regulates transcript profiles associated with early bipotent gonadal state and the first wave of pre-granulosa cells

We studied the transcriptomes of WT and *NR2F2*-KO COV434 cells using RNA sequencing (RNA-seq). Quality control analysis of RNA-seq data identified an outlier sample in the WT group, which is visualized in the principal component analysis (PCA) plot in Appendix Fig 3A. This sample was excluded in further analysis and a new PCA plot was generated, demonstrating that *NR2F2*-KO replicates (n=4) form a distinct cluster from WT replicates (n=3), which indicates that COUP-TFII deletion in COV434 cells resulted in significant transcriptional variation (Appendix Fig S3B).

*NR2F2*-KO cells had a total of 1724 differentially expressed genes (DEGs) when compared to WT COV434 cells. Of these, 1038 DEGs were downregulated (log2FC<-1) and 686 were upregulated (log2FC>1) (Fig 4A). Several DEGs were validated by RT-qPCR (Appendix Fig S4A,B). Gene markers selected from Guo et al. (Guo *et al*, 2021) and Garcia-Alonso et al. (Garcia-Alonso *et al*, 2022) to be associated with early somatic cells (*LHX9* and *VSNL1*), bipotent ESGCs (*RIMS4*), preGC-I (*FOXO1, SOX4*, and *STAT1*), supporting PAX8-expressing cells (*IGFBP3*), gonadal interstitium (*DCN*), and extragonadal mesonephros (*GATA2*) were downregulated in the *NR2F2*-KO COV434 cells (Fig 4B). Other cell markers of early supporting cells, such as *KITLG, SP3, AXIN2*, and *PAX8*, and other genes related to the bipotent state or ovary/testicular commitment, such as *GATA4, NR5A1, WT1, SOX9, FOXL2*, and *WNT4*, were not differentially expressed (Fig 4B,D). Gene ontology (GO) analysis for biological processes revealed that the most enriched term for genes downregulated in *NR2F2*-KO cells was “axon guidance” (Fig 4C), which is a transcriptional pattern of gonadal supporting *PAX8*-expressing cells (Garcia-Alonso *et al*, 2022). When considering the upregulated DEGs, the most enriched GO term was “angiogenesis”. Indeed, the expression of the endothelial cell marker *VEGFA* increased in *NR2F2*-KO COV434 cells (Fig 4B) and the upregulated DEGs were enriched in the cluster of perivascular and smooth muscle cells observed in the scRNA-seq datasets of developing human ovary and mesonephros (Appendix Fig S4C).

**Fig 4.**
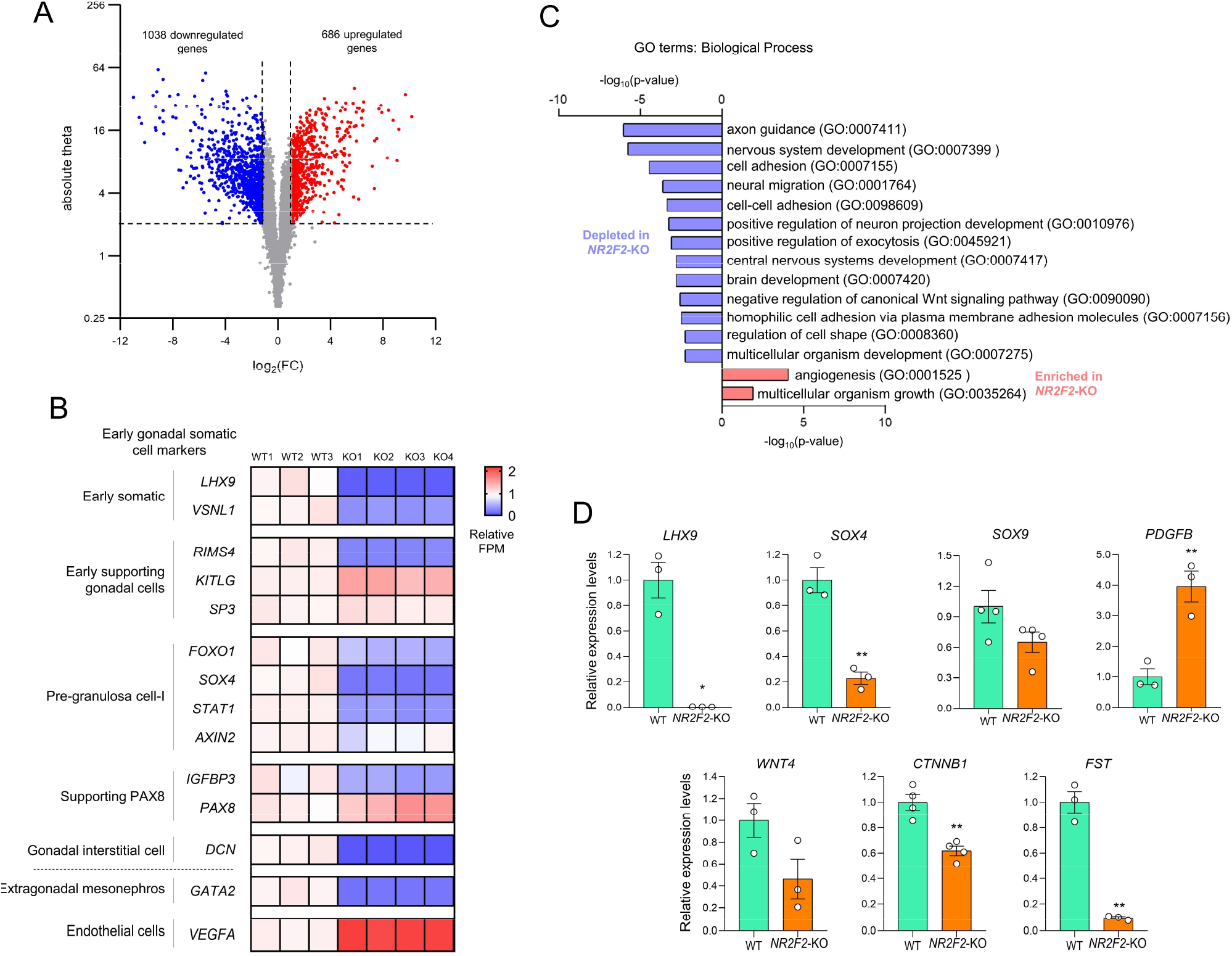
Transcriptome analysis of WT and the *NR2F2*-KO COV434 cells. (**A**) Volcano plot representing downregulated (log_2_FC <-1) and upregulated (log_2_FC >1) differentially expressed genes (DEGs) between *NR2F2*-KO and WT COV434 cells. Absolute theta >2 was used as a cutoff. (**B**) Heatmap representing gene expression levels of cell markers associated with early gonadal somatic cells, extragonadal mesonephros, and endothelial cells in the RNA-seq of WT and *NR2F2*-KO COV434 cells. The values of fragments per million mapped fragments (FPM) are expressed relative to the WT mean. (**C**) Gene ontology analysis of biological processes for downregulated and upregulated DEGs (*NR2F2*-KO *versus* WT COV434 cells). (D) RT-qPCR assay comparing relative mRNA expression of transcripts related to early gonadal somatic cells (*LHX9* and *SOX4*), testis (*SOX9* and *PDGFB*), and ovary development (*WNT4, CTNNB1*, and *FST*) between WT and *NR2F2*-KO COV434 cell clones. *S8* was used as a reference gene. Values are mean ± SEM (n=3-4). Student’s t-test with Welch’s correction, *p<0.05, **p<0.01. Source data are available online for this figure.

### Gene expression in NR2F2-KO COV434 cells indicates a loss of early supporting female gonadal fate and adoption of interstitial cell fate

During embryogenesis and organogenesis, COUP-TFII is known to regulate cell stemness and cell lineage commitment (Polvani *et al*, 2020). Thus, we assessed the cell type for which the gene signatures of WT and *NR2F2*-KO COV434 cells are enriched. We used the online platform WebCSEA (Dai *et al*, 2022) to compare the unique gene signatures of these cell clones with the gene sets of several cell types from human fetal organ systems. The unique gene signatures of WT and *NR2F2*-KO cells were determined by selecting the top 100 expressed genes (based on FPM) in each of these cells after excluding the overlapping 378 genes among their 500 top expressed genes, as schematized in Fig 5A. These overlapping genes are enriched for general biological processes, such as translation and protein folding, and could veil differences between the gene signatures (Appendix Fig S5). When compared with gene sets for the fetal reproductive system, the WT COV434 cells demonstrated a gene signature most resembling female gonadal epithelial progenitors, which give rise to the pre-granulosa cells (Fig 5B). Conversely, *NR2F2*-KO cells lost enrichment of the supporting signalling profile and acquired gene signatures more similar to gonadal interstitial fibroblasts and placental trophoblasts. *NR2F2*-KO COV434 cells did not acquire a pre-Sertoli transcriptional state. The unique gene signature of *NR2F2*-KO cells was also enriched in interstitial/stromal cells from other fetal organs (Fig 5B).

**Fig 5.**
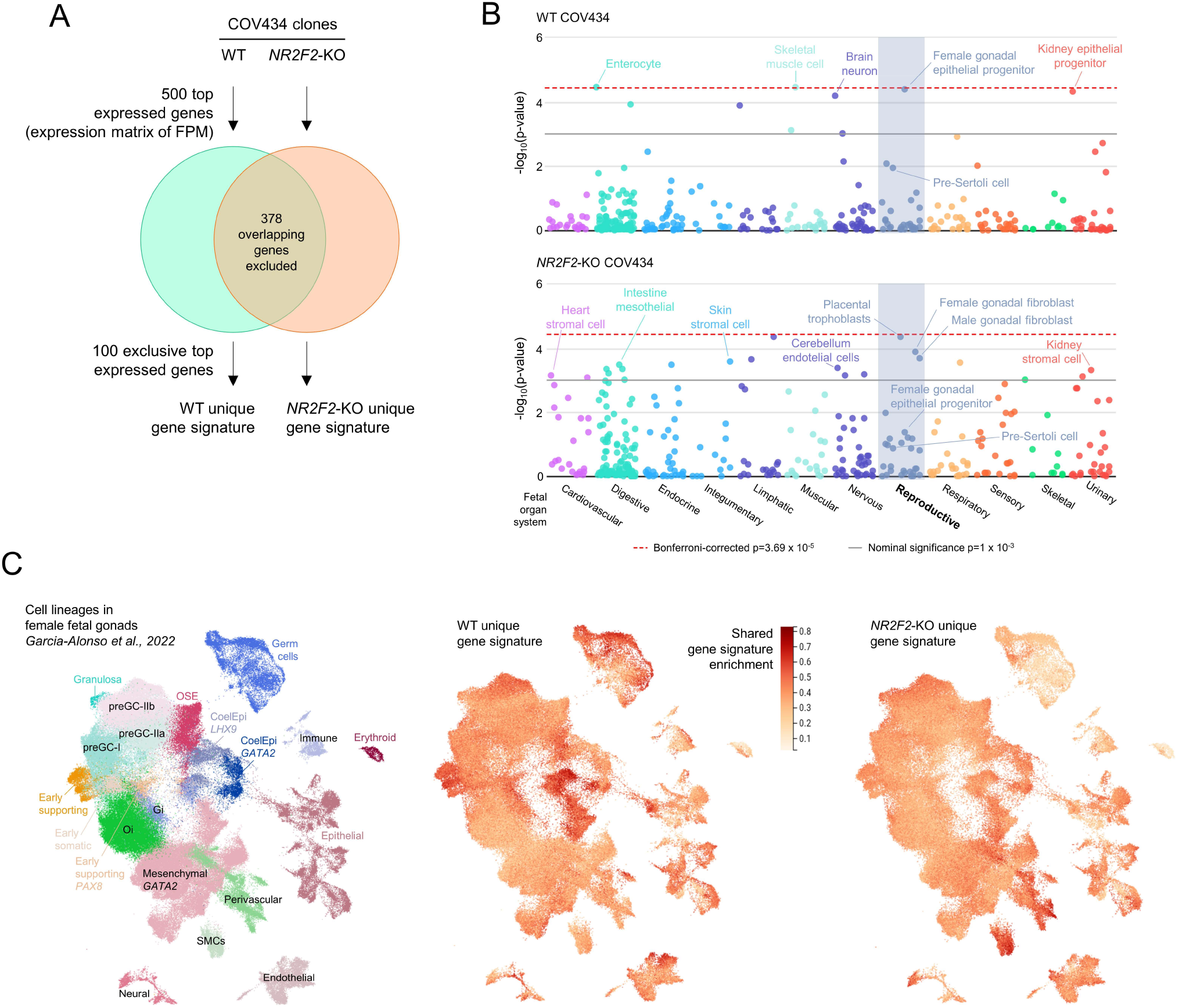
Comparison of the unique gene signatures of WT and the *NR2F2*-KO COV434 cells to the transcriptome of cell types from fetal organ systems and female fetal gonads. (**A**) Venn diagram depicting the adopted strategy to obtain the unique gene signatures for WT and *NR2F2*-KO COV434 cells from RNA-seq data. FPM, fragments per million mapped fragments. (**B**) The unique gene signatures for WT and *NR2F2*-KO cells were compared with tissue-cell type expression signatures of different human fetal organs using the online platform WebCSEA. Data show the -log_10_(p-value) generated for each query of the cell-type specificity enrichment analysis. Each dot represents one tissue-cell type characterized by scRNA-seq experiments. The red dashed line indicates the Bonferroni-corrected significance by 1355 tissue-cell types. The grey solid line indicates the nominal significance. (**C**) UMAP of cell lineages in the scRNA-seq datasets of developing ovary and mesonephros obtained from human female fetuses (Garcia-Alonso *et al*, 2022). The color scale represents the shared gene signature enrichment for WT and *NR2F2*-KO COV434 cells. CoelEpi, coelomic epithelium; OSE, ovarian surface epithelium; preGC, pre-granulosa cell; Gi, gonadal interstitial; Oi, ovarian interstitial; SMC, smooth muscle cell. Source data are available online for this figure.

To align WT and *NR2F2*-KO gene signatures to data from human fetal gonads, we used scRNA-seq datasets of developing ovary and mesonephros from human female fetuses (Garcia-Alonso *et al*, 2022). When focused on the gonadal epithelial progenitor cells, the gene signature from WT COV434 cells was enriched in subpopulations of *LHX9*-expressing coelomic epithelial cells and early supporting cells. We also observed enrichment of mesenchymal cells of the mesonephros (expressing *GATA2*) and germ cells. However, *NR2F2*-KO cells lost enrichment of the epithelial progenitor cell lineages and more closely resembled the gene signatures of the interstitial smooth muscle and perivascular cells (Fig 5C). Indeed, the most expressed gene in the *NR2F2*-KO unique gene signature was *RGS5* (Source data for Fig 4 and 5), which has been previously identified as a smooth muscle cell marker in the developing gonads (Guo *et al*, 2021).

## Discussion

This is the first study to address the genetic network regulated by *NR2F2*/COUP-TFII in a supporting gonadal cell context and to investigate *NR2F2* expression during human bipotential gonad commitment. By analyzing scRNA-seq datasets of somatic cell lineages from fetal gonads and by differentiating iPSCs into bipotential gonad-like cells *in vitro*, we demonstrated that the human *NR2F2* is highly upregulated during bipotential gonad development, being detected in early somatic cells that precede the steroidogenic cell emergence in the undifferentiated gonad. The generation of the granulosa-like cell COV434 *NR2F2*-KO suggested that COUP-TFII regulates pathways involved in the early bipotential gonad and the first wave of pre-granulosa cells. The identification of loss-of-function genetic variants in *NR2F2* in individuals with *SRY*-negative 46,XX T/OT-DSD (Carvalheira *et al*, 2019; Bashamboo *et al*, 2018) has suggested that COUP-TFII is a pro-ovary factor during sex development. Altogether, our data corroborate the hypothesis that *NR2F2*/COUP-TFII is involved in this pro-ovary module and that loss-of-function variants in the *NR2F2* gene anticipate the multipotent state disruption of ESGCs, leading to testicular or ovotesticular development in 46,XX individuals (Fig 6).

**Fig 6.**
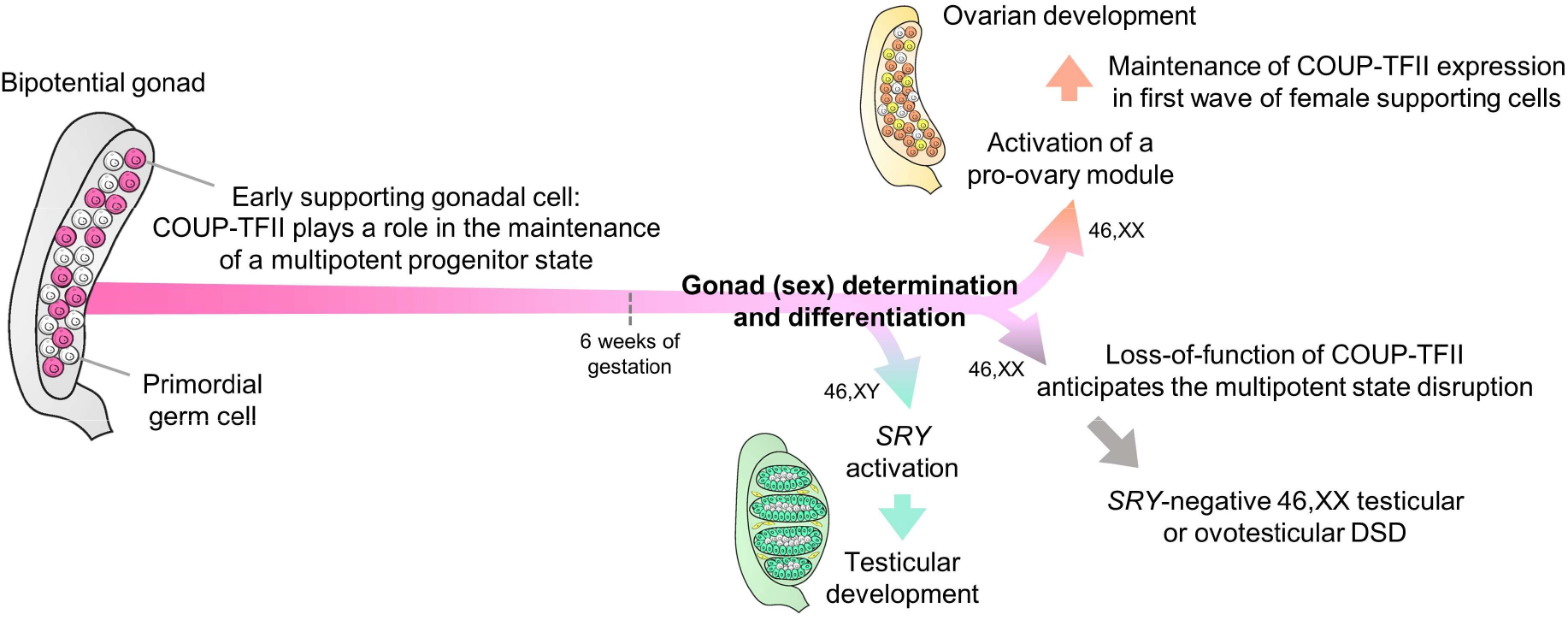
Proposed model for COUP-TFII function during early gonadal development in humans. DSD, differences of sex development.

Analysis of scRNA-seq datasets revealed that ESGCs expressing the *SRY* gene, thus differentiating into fetal Sertoli cells, lost *NR2F2* expression, which became restricted to interstitial cells after testis differentiation. We also found that *NR2F2* transcript abundance is much higher in granulosa-like COV434 cells than in Sertoli-like NT2/D1 cells, supporting this sex-biased expression pattern. Recently, Pierson Smela et al. (Pierson Smela *et al*, 2023) showed that the transcriptomes of two granulosa-like cell lines, COV434 and KGN, both have modest similarity to the fetal ovary, including the enrichment for *NR2F2* expression. This data justified our use of the COV434 cell line to study COUP-TFII function.

We examined the expression of the four *NR2F2* transcript variants individually in a human gonadal context. Previous studies showed that the action of the canonical COUP-TFII isoform A could be enhanced or inhibited by the other isoforms in a manner dependent on the genomic context and the cellular type (Rosa & Brivanlou, 2011; Yamazaki *et al*, 2013; Davalos *et al*, 2023), emphasizing the importance to study all COUP-TFII isoforms. *NR2F2* frameshift variants reported by Bashamboo et al. (Bashamboo *et al*, 2018) in *SRY*-negative 46,XX T/OT-DSD patients are predicted to affect the COUP-TFII isoform A only. Indeed, our data suggest that isoform A is the major COUP-TFII isoform expressed during human gonad development since transcript v1, encoding isoform A, is nearly 1000-fold more highly expressed than other *NR2F2* variants in bipotential gonad-like cells derived from iPSCs. Additionally, *in vitro* differentiation of bipotential gonad-like cells demonstrated that *NR2F2* is upregulated along with bipotential gonad markers, such as *GATA4*.

COUP-TFII and GATA4 have been shown to physically and functionally cooperate in the regulation of gene expression in murine Leydig cells (Mehanovic *et al*, 2022).

RNA-seq analysis of *NR2F2*-KO COV434 cells has also highlighted additional pathways in which COUP-TFII may act. The most enriched GO term for downregulated genes in these KO cells was “axon guidance”. Interestingly, Stévant et al. (Stévant *et al*, 2018) found “axon development” as the most enriched term for the upregulated genes in early progenitor cells from male murine gonads. Garcia-Alonso et al. (Garcia-Alonso *et al*, 2022) reported that supporting *PAX8*-expressing cells located at the gonadal–mesonephric interface during gonad development show a transcriptional pattern of axon guidance factors, which is associated with stem cell maintenance and renewal. Indeed, COUP-TFII is an important regulator of cellular fate choice, being implicated in the regulation of the mesenchymal stem cell state by modulating the WNT signaling, RUNX2 activity, as well as *Pparg* and *Sox9* expression (Polvani *et al*, 2020),(Xie *et al*, 2022). These findings together with our data showing that *NR2F2* is expressed in early gonadal cell populations retaining a multipotent state suggest that COUP-TFII may regulate cell stemness and lineage commitment during gonadal development.

The early gonadal somatic cells are multipotent progenitors which can give rise to either supporting (pre-granulosa or fetal Sertoli cells) or interstitial cell lineages (Stévant *et al*, 2019; Guo *et al*, 2021). Studies with transgenic *Sry* reporter mice showed that female ESGCs can activate the *Sry* promoter in the same time window as male ESGCs (Albrecht & Eicher, 2001; Harikae *et al*, 2013). Additionally, human pre-granulosa cells express *SOX9* in an early stage of their differentiation (Li *et al*, 2017), which may be associated with the previously reported SF1-mediated upregulation of *SOX9* expression in an SRY-independent manner (Ming *et al*, 2022). Indeed, female and male ESGCs display similar transcriptomic programs harboring the expression of factors predicted as pro-testicular, such as *NR5A1, WT1*, and *GATA4*, which participate in the activation of *SRY* expression in XY embryos (Mäkelä *et al*, 2019; Reyes *et al*, 2023). However, the pro-testicular signaling pathway might be balanced by a still not fully understood pro-ovarian pathway even in the early population of gonadal uncommitted somatic cells. The mutual neutralization of these antagonistic pathways maintains a multipotent state. *SRY* activation disrupts this balance, leading to testis development in XY embryos; however, the impaired function of pro-ovarian factors may also disrupt the transcriptional plasticity of ESGCs and instead drive them into commitment to the testicular pathway, leading to Sertoli cell development in XX embryos (Fig 6). We propose that COUP-TFII is a member of the pro-ovary pathway counteracting testicular commitment in the early somatic cells, being downregulated during *SRY* activation.

The transient bipotential state of early supporting cells is temporally asymmetric between testis and ovary development. In contrast to fetal Sertoli cells, pre-granulosa cells maintain the expression of stem cell-related genes, remaining in a progenitor-like state for several days (Stévant *et al*, 2019). By comparing the unique gene signatures of WT and *NR2F2*-KO COV434 cells, we demonstrated that *NR2F2*-KO cells lost the enrichment for female-supporting gonadal progenitor and acquired gene signatures more similar to gonadal interstitial cells, such as smooth muscle and perivascular cells, which is in line with the enrichment of the GO term “angiogenesis” for the upregulated genes in *NR2F2*-KO cells. Therefore, we propose that COUP-TFII has a role in the maintenance of a multipotent progenitor state in ESGCs and pGC-I, where its role is to repress commitment into supporting cells during the time window in which male and female ESGCs are prone to activate pro-testicular factors. The absence of a pro-testicular context in the COV434 cells may explain why we observed the enrichment of an interstitial transcriptional pattern instead of a fetal Sertoli state upon COUP-TFII ablation.

Several studies, mostly in Leydig cells, have shown a competition between COUP-TFII and SF1 for binding to overlapping response elements in the promoter region of genes encoding steroidogenic enzymes (Hattori & Fukami, 2023). *NR2F2* expression is detected during the commitment of gonadal interstitial cells to steroidogenic cells and in fetal Leydig cells; however, it is progressively repressed during adult Leydig cell differentiation, allowing the SF1-mediated upregulation of steroidogenic enzymes (van den Driesche *et al*, 2012; Guo *et al*, 2021). The inverse expression of *NR2F2* and steroidogenic genes may reflect the role of COUP-TFII in preserving the progenitor Leydig cell pool by repressing their maturation via antagonizing SF1 (Bhattacharya & Dey, 2023). COUP-TFII and SF1 physically interact in mouse Leydig cells and display antagonistic regulation patterns in the promoter of *Nr0b1* (DAX1), which also participates in gonadal development (Yu *et al*, 1998; Di-Luoffo *et al*, 2022). Indeed, COUP-TFII co-binds with different nuclear receptors in specific DNA motifs for cooperation or competition (Boudot *et al*, 2011). However, a potential interplay between COUP-TFII and SF1 as a pro-ovary and a pro-testis factor, respectively, during supporting cell commitment is yet to be investigated.

Using scRNA-seq on ovarian and testicular *Nr5a1*-expressing somatic cells during murine sex determination, Stévant et al. (Stévant *et al*, 2019, 2018) demonstrated that, while a pool of somatic progenitors differentiates into supporting cells, the remaining interstitial progenitors gradually undergo transcriptional changes restricting their competence toward a steroidogenic fate but retaining their multipotent state. These cells, which maintain *Nr2f2* expression, can be trans-differentiated into supporting cells (Zhang *et al*, 2015). Another open question is if COUP-TFII could also participate in the maintenance of this multipotent state during the early development of interstitial and steroidogenic cells, providing a source of cells still able to shift their differentiation into the supporting lineage.

COV434 is an adult tumor cell line and its genetic background poses a significant limitation in understanding fetal gonadal development. The COV434 cells are negative for the granulosa cell marker FOXL2 and Karnezis et al. (Karnezis *et al*, 2021) recently suggested that the original tumor that originated this cell line has a histologic identity as a small cell carcinoma of the ovary. Therefore, a robust human model that recapitulates *in vivo* ovarian development is urgently required to advance the knowledge regarding the pro-ovary module participating in the early gonadal commitment during human gonad development. Additionally, *NR2F2*/COUP-TFII and *NR2F1*/COUP-TFI show a high degree of evolutionary conservation, suggesting redundancy and overlapping functions (Polvani *et al*, 2020). This homologous transcription factor is expressed during the early gonadal development in mice and humans, although in a lower abundance than *NR2F2* (Stévant *et al*, 2019; Guo *et al*, 2021). To date, no genetic variants have been identified in the *NR2F1* gene in patients with DSD. Further studies are needed to understand if *NR2F1*/COUP-TFI plays a role in gonadal development.

In summary our data suggest that COUP-TFII is a member of the pro-ovary pathway counteracting the commitment of early supporting gonadal cells to Sertoli cells and maintaining a multipotent state necessary until the later commitment to pre-granulosa cell differentiation occurs, thus regulating the temporal asymmetry between testis and ovary development (Fig 6). Therefore, we propose that COUP-TFII plays dual roles during gonad development, regulating both the supporting cell fate and interstitial cell differentiation.

## Materials and Methods

### Cell culture of COV434, NT2/D1, and HepG2 cell lines

The human granulosa-like cell line COV434 (ECACC, #07071909), the human Sertoli-like cell line NT2/D1 (NTERA-2 cl.D1 [NT2/D1]; Banco de Células do Rio de Janeiro, #0303), and the human hepatocyte cell line HepG2 (Banco de Células do Rio de Janeiro, #0103) were cultured in DMEM high glucose (Thermo Fisher Scientific) supplemented with 4 mM L-glutamine, 1.7 g/L sodium bicarbonate, 10% fetal bovine serum, 100 U/mL penicillin-streptomycin, and 0.5 μg/mL amphotericin B. The cell culture medium for NT2/D1 was also supplemented with 1 mM sodium pyruvate. Cells were kept in a humidified incubator at 37°C with 5 % CO_2_/95 % air.

### Single guide RNA design and vector construction

To design guide RNAs (gRNA) for the *NR2F2* gene, we used the program CRISPOR (Concordet & Haeussler, 2018) available online on http://crispor.tefor.net/. sgRNA sequences used are listed in Appendix Table S1. The two strands of the gRNA were annealed and cloned downstream of the human U6 promoter using the *Bbs*I (NEB) restriction site in the plasmid pU6-(BbsI)_CBh-Cas9-T2A-BFP (Addgene). The protocols for gRNA cloning and transformation of competent *E. coli* were performed as previously described (Ran *et al*, 2013). Sanger sequencing was applied to confirm that the vectors were correctly constructed.

### Flow-cytometry sorting and single-cell clone genotyping

COV434 cells were transfected with the plasmid cloned with the gRNA directed toward the 5’ region of exon 2 of the *NR2F2* gene using Lipofectamine 3000 (Thermo Fisher Scientific). Blue fluorescence protein (BFP)-positive cells were observed and photographed using a fluorescence microscope (Zeiss) 24 h after transfection. 48 h after transfection, cells were dissociated using Trypsin-EDTA and BFP-positive cells were individually seeded into wells of a 96-well plate for single-cell culture by fluorescence-activated cell sorting (FACS) using BD FACSAria III (BD Biosciences). As described by Zhang et al. (Zhang *et al*, 2020), single cells were kept on conditioned culture medium (medium from log-phase cells filtered through a 0.22 μm pore size filter supplemented with fresh cell culture medium 1:1). Medium was changed every 2 days. From day 17 after transfection, single cell-derived clones were observed under the phase contrast light microscope. Mycoplasma testing was performed by PCR and all samples tested negative.

Genomic DNA was extracted from cell clones using an in-house method as previously described (Kizys *et al*, 2012). PCR amplification of exon 2 of the *NR2F2* gene was performed using PCR Master Mix (Promega; primer pair is listed in Appendix Table S1). The amplicon was cloned into the TOPO™ vector using TOPO™ TA Cloning™ Kit (Thermo Fisher Scientific), which was used to transform competent *E. coli* for Sanger sequencing of individual alleles using Big DyeTM Terminator Cycle Sequencing Ready Reaction Kit in ABI Prism 3130xl Genetic Analyzer (Applied Biosystems) (Kizys *et al*, 2012). Genotyping array was performed using Infinium Global Screening Array -24 v3.0 (Illumina) to evaluate genome integrity.

### Protein extraction and western blot

COV434 cell clones and NT2/D1 cells were homogenized in RIPA lysis buffer (50 mM Tris, pH 7.5; 150 mM NaCl, 1% Nonidet P-40; 0.5% sodium deoxycholate; 1 mM EDTA and 0.1% SDS) supplemented with protease inhibitors (Protease Inhibitor Tablets, Thermo Scientific Pierce™) using Polytron® equipment (KINEMATIC). Western blotting was performed as previously described (Serrano-Nascimento *et al*, 2018). Briefly, proteins were blotted onto nitrocellulose membranes, blocked with 3% BSA solution, and incubated with the primary antibodies anti-COUP-TFII (1:1000, Abcam, #41859) and anti-Alpha-tubulin (1:2000, Sigma, #T9026). Membranes were then incubated with the corresponding secondary antibody conjugated to horseradish peroxidase. Blots were developed using the enhanced chemiluminescence (ECL) kit (Bio-Rad).

### Real-time quantitative PCR for COV434, NT2/D1, and HepG2 cells

RNA was extracted using RNeasy Plus Kit (QIAGEN). Reverse transcriptase reactions were performed with oligo(dT)_18_ using the M-MLV Reverse Transcriptase kit (Thermo Fisher Scientific). The cDNA samples were assayed in real-time quantitative PCR (qPCR) using the kit PowerTrack SYBR Green Master Mix (Thermo Fisher Scientific) at the thermocycler ABI PRISM 7500 Sequence Detection System (Applied Biosystems). The primer pairs were designed using the program NCBI/Primer-BLAST (Ye *et al*, 2012) and are indicated in Appendix Table S1. The expression of target genes was normalized using the reference gene *S8* (ΔCt) and represented as a percentage of *S8* (2^-ΔCt^) or relatively to a control/reference group (2^-ΔΔCt^) (Schmittgen & Livak, 2008).

### H&E and fluorescent immunocytochemistry

COV434 cell clones cultured on coverslips were fixed in 4% paraformaldehyde, stained with hematoxylin and eosin (H&E) solution, and mounted on glass slides for histological analysis. Fixed cells were also used for fluorescent immunocytochemistry studies. Cells were incubated with blocking solution (0.1% Triton X-100 and 2% BSA in PBS) for 30 minutes at room temperature and then overnight at 4°C in blocking solution containing mouse anti-Alpha-tubulin (1:500, Sigma, #T9026). Next, cells were incubated for 1 h at room temperature with anti-mouse secondary antibody conjugated to Alexa Fluor 594 and then for 1 h at room temperature with Phalloidin-FITC 488 (1:500, Sigma, #49409). DAPI (4,6-diamidino-2-phenylindole) was used for nuclear identification. Negative controls were performed in the absence of primary antibody. Fluorescence images were acquired under a Nikon E800 microscope.

### RNA-Sequencing and data analysis

RNA was extracted from four replicates of WT and *NR2F2*-KO COV434 cells using RNeasy Plus Kit (QIAGEN). The RNA quality and quantity were assayed on NanoDrop spectrophotometer and Qubit fluorometer (Thermo Fisher Scientific). RNA libraries were constructed using Zymo-Seq RiboFree Total RNA Library Kit (Zymo Research) following the manufacturer’s protocol. Each library was analyzed on Agilent Bioanalyzer (Agilent Technologies) and by qPCR for quality and quantification assessment and sequenced on NovaSeq 6000 in a flow-cell SP 2×150pb (Illumina).

Quality control with FastQC (Andrews, 2010) and MultiQC (Ewels *et al*, 2016) was done in all samples before alignment, after alignment and after gene counting. Samples were aligned to the human reference by using STAR (Dobin *et al*, 2013), using Hg38 and GTF v. 103 from the Ensembl (Herrero *et al*, 2016) project. Genes were counted with Rsubread (Liao *et al*, 2019) program and the featurecounts function. Gene counts were normalized using the fragments per million mapped fragments (FPM) function of DeSeq2 (Love *et al*, 2014). Then, low expression genes were filtered off using Noiseq (Tarazona *et al*, 2015) filter method 3. The differential expression (DE) was done using the NoiseqBio (Tarazona *et al*, 2015) function. DAVID (Huang *et al*, 2009; Sherman *et al*, 2022) was used to calculate gene ontology enrichment for significantly upregulated (log2FC >1, absolute theta >2) and downregulated (log2FC <−1, absolute theta >2) genes. The gene signatures of WT and *NR2F2*-KO COV434 cells were determined by selecting the unique most expressed genes based on FPM values, as schematized in Fig 5A.

### Analysis of single-cell RNA sequencing datasets of human fetal tissues

*NR2F2* expression was studied in single-cell RNA sequencing (scRNA-seq) datasets of somatic cell lineages from male and female human gonads between 6- and 21-weeks gestation available online on the Reproductive Cell Atlas (https://www.reproductivecellatlas.org/) (Garcia-Alonso *et al*, 2022). Gene lists representing the gene signatures of WT and *NR2F2*-KO COV434 cells, and the up and downregulated genes (*NR2F2*-KO *versus* WT) were uploaded into the online platforms Reproductive Cell Atlas and WebCSEA (https://bioinfo.uth.edu/webcsea/) (Dai *et al*, 2022) for comparison with the gene sets of several cell types from human fetal organ systems. The scRNA-seq data for reproductive organs available in the WebCSEA are from two female fetal gonads (11- and 26-weeks gestation) and two male fetal gonads (11- and 12-weeks gestation) (GEO: Series GSE134355).

### Human iPSC culture, monolayer differentiation, and real-time quantitative PCR

The differentiation experiments were performed as previously described (Knarston *et al*, 2020). Briefly, the human induced pluripotent stem cell (iPSC) lines CRL1502 (female) (generated by E.J. Wolvetang, The University of Queensland, Australia) and PCS_201 (male) (American Type Culture Collection, USA) were expanded in Essential 8 medium (E8; Thermo Fisher Scientific). One day prior to differentiation, cells were plated at 10,000 cells/cm^2^ on Vitronectin (STEMCELL Technologies). On day 0, the medium was replaced with Essential 6 medium (E6). Cells were cultured for 4 days with 3 μM CHIR (R&D Systems); 3 days with 200 ng/mL FGF9 (R&D Systems), 1 μg/mL heparin (Sigma-Aldrich), and 10 ng/mL BMP4 (R&D Systems); and 2 days without growth factors. Medium was changed every 2 days. RNA was harvested at days 0, 4, 7, and 9.

The qPCR assays were performed as described by Knarston et al. (Knarston *et al*, 2020). Briefly, RNA was extracted using the ReliaPrep RNA Cell Miniprep System (Promega) and cDNA was synthesized using the GoScript reverse transcriptase system (Promega). qRT-PCR was performed with GoTaq qPCR Master Mix (Promega) on the LightCycler480 (Roche). Primer sequences were described previously (Knarston *et al*, 2020) or can be found in Appendix Table S1. The expression of target genes was represented by normalizing with the expression of the reference gene *GAPDH* (2^-ΔCt^) (Schmittgen & Livak, 2008).

### Statistical analysis

GraphPad Prism software version 8.0 (GraphPad Software Inc.) was used for all statistical analyses. Data were tested for normality with D’Agostino and Pearson tests. For two-group comparisons, the two-tailed Student’s t-test was used with Welch’s correction when applicable. For comparisons between multiple groups, the one-way ANOVA test was followed by a Tukey test. Data were expressed as mean ± SEM. P < 0.05 was considered statistically significant.

## Acknowledgements

We would like to thank the Laboratory of Molecular and Translational Endocrinology (LEMT/UNIFESP) for their technical support, and Dr. Rui Maciel for laboratory support. We would also like to thank the Conselho Nacional de Desenvolvimento Científico e Tecnológico (CNPq), the Coordenação de Aperfeiçoamento de Pessoal de Nível Superior (Capes) and the São Paulo Research Foundation (FAPESP) for their financial support, including the sponsorship FAPESP #2022/10804-9 to Marina M. L. Kizys. This work was supported by a grant from FAPESP (#2021/00684-3) and a grant from CNPq (#200262/2022-0) to Magnus R. Dias da Silva.

## Author contributions

Lucas G. A. Ferreira: Conceptualization; formal analysis; validation; investigation; visualization; methodology; writing – original draft. Marina M. L. Kizys: Conceptualization; methodology; project administration. Gabriel A. C. Gama: Data curation; software; formal analysis; methodology. Svenja Pachernegg: Formal analysis; methodology; supervision; investigation; writing – review and editing. Gorjana Robevska: Methodology; investigation. Andrew H. Sinclair: Resources; supervision; writing – review and editing. Katie L. Ayers: Resources; investigation; supervision; writing – review and editing. Magnus R. Dias da Silva: Conceptualization; project administration; supervision; resources; funding acquisition.

## Data availability

The RNA-Seq data produced in this study was submitted to SRA (SUB13718909) and GEO on August 4, 2023 under the GEO project PRJNA982097 (NCBI tracking system #24215611).

